# A dueling-competent signal-sensing module guides precise delivery of cargo proteins into target cells by engineered *Pseudomonas aeruginosa*

**DOI:** 10.1101/2022.09.02.506313

**Authors:** Li-Li Wu, Shuangquan Yan, Tong-Tong Pei, Ming-Xuan Tang, Hao Li, Xiaoye Liang, Shuyang Sun, Tao Dong

## Abstract

To recognize and manipulate a specific microbe of a crowded community is a highly challenging task in synthetic biology. Here, we introduce a highly-selective protein delivery platform, termed DUEC, which responds to direct contact of attacking cells by engineering the tit-for-tat/dueling response of H1-T6SS (type VI secretion system) in *Pseudomonas aeruginosa*. Using a Cre-recombinase-dependent reporter, we screened H1-T6SS secreted substrates and developed Tse6^N^ as the most effective secretion tag for Cre delivery. DUEC cells can discriminately deliver the Tse6^N^-Cre cargo into the cytosol of T6SS^+^ but not T6SS^−^ *Vibrio cholerae* cells in a mixed population. These data demonstrate that the DUEC cell is not only a prototypical physical-contact sensor and delivery platform but also may be coupled with recombination-based circuits with the potential for complex tasks in mixed microbial communities.

## Introduction

Microbes exist in complex communities and collectively contribute to their diverse functions in the natural and host environments. The microbial community can be modulated by antibiotics, probiotics, and nutrients on a global scale. However, tools for changing a select microbe within a complex community remain scarce. *Pseudomonas aeruginosa*, an important pathogen, can sense and remember the precise location of physical contact of a neighboring bacterium and carry out a retaliatory response termed dueling between sister cells or tit-for-tat against heterologous attackers ^1–3^. Such unique response is mediated by the type VI protein secretion system (T6SS) in *P. aeruginosa*, and may be used to selectively modulate a microbe within a complex microbial community. However, the underlying mechanism is not fully understood, hindering its use as a signal-sensing module for detecting cell-perturbing direct-contact signals.

T6SS is a double-tubular contractile weapon commonly employed by gram-negative bacteria to secrete toxic effectors and kill competitors in diverse environments upon direct contact ^4–6^. The outer sheath contracts and releases a substantial amount of energy that drives the inner tube and its associated spike complex and effectors deep into the cytosol of neighboring cells including gram-negative and gram-positive bacteria and fungi ^7–12^. Sheath length likely dictates the working distance of T6SS given that delivery into a target cell is mostly contact-dependent with few exceptions ^13–17^. Notably, receiving cells are not killed by penetration of the incoming tube but by the co-delivered effectors ^9,18,19^. These effectors may not only dictate functions but also directly contribute to T6SS assembly because deletion of multiple effector genes abolishes T6SS assembly in several species ^9,18,20^. Similar to the spike complex proteins, VgrG and PAAR, some effectors can also serve as carriers when fused with cargo proteins ^8,9,21,22^. Therefore, rather than deleting effector genes, an effective strategy has been developed in *Vibrio cholerae* and *Aeromonas dhakensis* to use catalytic mutations of effectors to detoxify the T6SS while maintaining the delivery function ^9,20,22^. However, it is challenging to apply such strategy in much more complex systems such as the T6SS in *P. aeruginosa*, which has three independent T6SS gene clusters (H1-, H2-, and H3-T6SS), each secreting a specific set of effectors ^5,23–28^. Eight effectors (Tse1-8) of H1-T6SS with different modes of secretion have been identified ^13,29–34^ but whether they contribute to T6SS assembly and can serve as cargo carriers has not been explored.

Tools for specifically responding to a physical contact signal are currently lacking in the vast pool of synthetic biology modules ^35–38^. The tit-for-tat/dueling response may fill the gap and can be used to manipulate specific microbes among mixed communities. This response is controlled by the threonine phosphorylation pathway (TPP) comprising a cascade of signal-sensing and output proteins, including the membrane-associated TagQRST sensor, the PpkA kinase that phosphorylates Fha1 of the H1-T6SS for activating H1-T6SS assembly, and the PppA phosphatase that carries out dephosphorylation of Fha1 for inactivation ^3,39–41^. Upon sensing membrane-perturbing signals from the T6SS of a heterologous attacker or a sister cell, the RP4 conjugation, or extracellular DNA, this pathway activates the H1-T6SS and, in the case between sister cells, initiates a counterattack within seconds precisely at the attacking location ^1–3,42^. However, heterologous expression of the TPP as a conventional signal sensing module is challenging due to its multi-component complexity, specific protein-interaction phosphorelay, and unresolved mechanism.

To address this challenge, we report an alternative strategy by engineering the *P. aeruginosa* H1-T6SS effectors so that the TPP can be employed *in situ* to control the delivery of cargo proteins and coupled with a Cre-mediated genetic switch. The resultant strain DUEC (<u>due</u>ling <u>c</u>ompetent) has lost its antibacterial activities but could differentiate T6SS^+^ and T6SS^−^ *V. cholerae* and selectively deliver Cre recombinase into T6SS^+^ *V. cholerae* cells to induce recombination. The DUEC system could thus function as a direct-contact response module for precise protein delivery to specific cells in mixed communities.

## Results

### Systematic inactivation reveals effector activities are dispensable for the H1-T6SS activities

To employ the TPP-controlled dueling as a synthetic biology module, we postulate that the strong antibacterial activity of the H1-T6SS may be an unwanted feature for a general delivery tool. To detoxify the H1-T6SS, we systematically introduced chromosomal mutations to the catalytic sites of known effectors except that the gene encoding Tse5 was deleted because of its unknown toxicity. The resultant mutant, lacking all eight effector activities, was named DUEC hereafter for simplicity. Time-lapse fluorescence microscopy analysis shows that DUEC cells exhibited active H1-T6SS assembly and dueling that are comparable to wild-type cells (Fig. 1A-1B). In addition, when using the T6SS-active *V. cholerae* strain V52 as a competitor, we found that DUEC cells exhibited abolished antibacterial activities similar to the T6SS deletion mutant (Fig. 1C). These results indicate that the H1-T6SS assembly and dueling do not require effector activities.

**Fig. 1.**
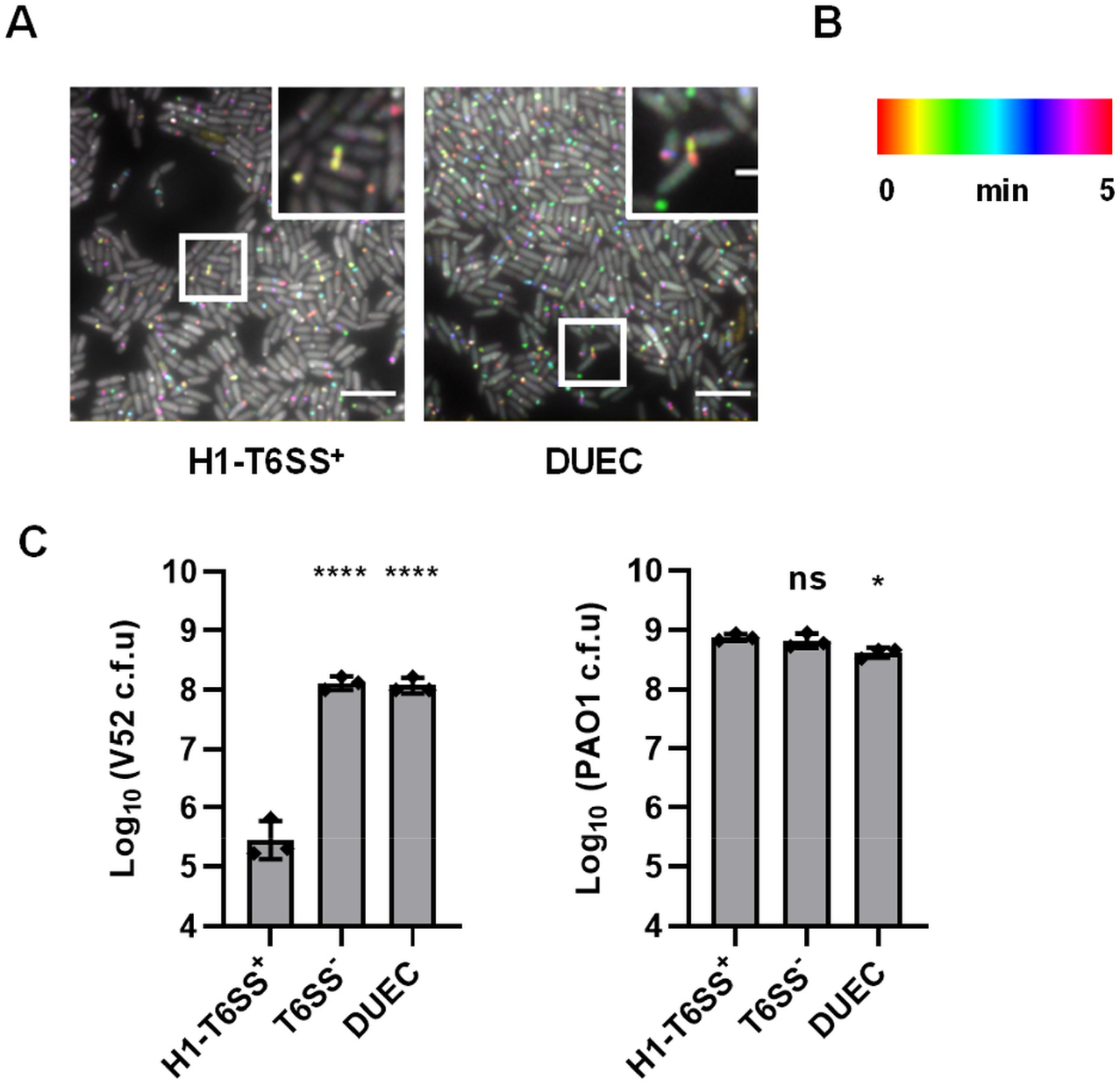
DUEC is T6SS-dueling competent but defective in bacterial killing. A, Time-lapse imaging of TssB1-sfGFP signal captured every 10 s for 5 min and temporally color-coded in T6SS^+^ mutant (Δ*retS*, Δ*tssB2*, Δ*tssB3*, TssB1-sfGFP) and eight-effector-inactivated mutant DUEC. A representative image of 30 μm × 30 μm field of cells with a 2 × magnified 5 μm × 5 μm inset (marked with a box) of a selected region is shown. Scale bar is 5 μm for the large field of view and 1 μm for the insets view and applied to all. B, Color scale used to temporally color code the TssB1-sfGFP signal. C, Survival of *V. cholerae* V52 and *P. aeruginosa* PAO1 after competition assay. Genotypes of PAO1 are indicated at the bottom. T6SS^−^: Δ*retS*, Δ*tssB1*, Δ*tssB2*, Δ*tssB3*. Killer and prey cells were mixed at the ratio of 5: 1. Error bars indicate the mean ± standard deviation of at least three different biological duplicates. One-way ANOVA with Dunnett’s multiple comparisons test compared with H1-T6SS^+^ mutant. \**P* <0.05, *\*\*\*\*P* < 0.0001; ns, not significant. Source data are provided as a Source Data file.

### H1-T6SS delivers Cre as a fused cargo to VgrG proteins

Unlike conventional signal transduction modules that contain a transcriptional regulator for downstream output, the outcome of the TPP pathway is the assembly and firing of the H1-T6SS. We postulated that an effective way to transform the T6SS secretion event into a genetically amendable module is to couple with the secretion of the Cre recombinase which could specifically recognize DNA sequences containing two *loxP* sites and induce recombination. We have recently built a pFIGR reporter system featuring a *loxP*-flanked ampicillin resistance cassette (Amp) that blocks the translation of a downstream gentamicin gene (Gen). The inhibition can be relieved by Cre-mediated recombination resulting in the loss of Amp resistance and the gain of Gen resistance. Taking advantage of this reporter, we set out to determine how to secrete Cre by the H1-T6SS.

Because VgrG proteins form a spike complex sitting at the tip of the inner tube and can carry both natural and engineered effector domains^21,43^, we hypothesized that adding Cre as a C-terminal fusion to the three H1-T6SS VgrG proteins (VgrG1a/1b/1c) may be an effective delivery approach. Thus, we constructed VgrG-Cre fusions, respectively (Fig. 2A). Using *V. cholerae* strains carrying the pFIGR reporter, we tested the recombination efficiency of donor strains expressing different VgrG-Cre fusions and used Cre-only vectors and the H1-T6SS null *tssB*1 deletion donor as negative controls. By co-incubating donor and recipient cells, we found that VgrG1a produced a 10-fold higher number of Gen^R^ recombinants than the negative controls and the other two VgrG fusions (Fig. 2A and Fig. S1A-S1C). These results suggest that the increased Gen^R^ resistance was specifically due to H1-T6SS mediated delivery of VgrG1a-Cre while the background levels of Gen^R^ mutant in the negative control samples might result from spontaneous mutations or recombination. Interestingly, we noticed that induction of VgrG1b-Cre and VgrG1c-Cre reduced donor cell survival, which may account, at least in part, for the ineffective delivery. Overall, these data indicate that Cre can be delivered by the H1-T6SS.

**Fig. 2.**
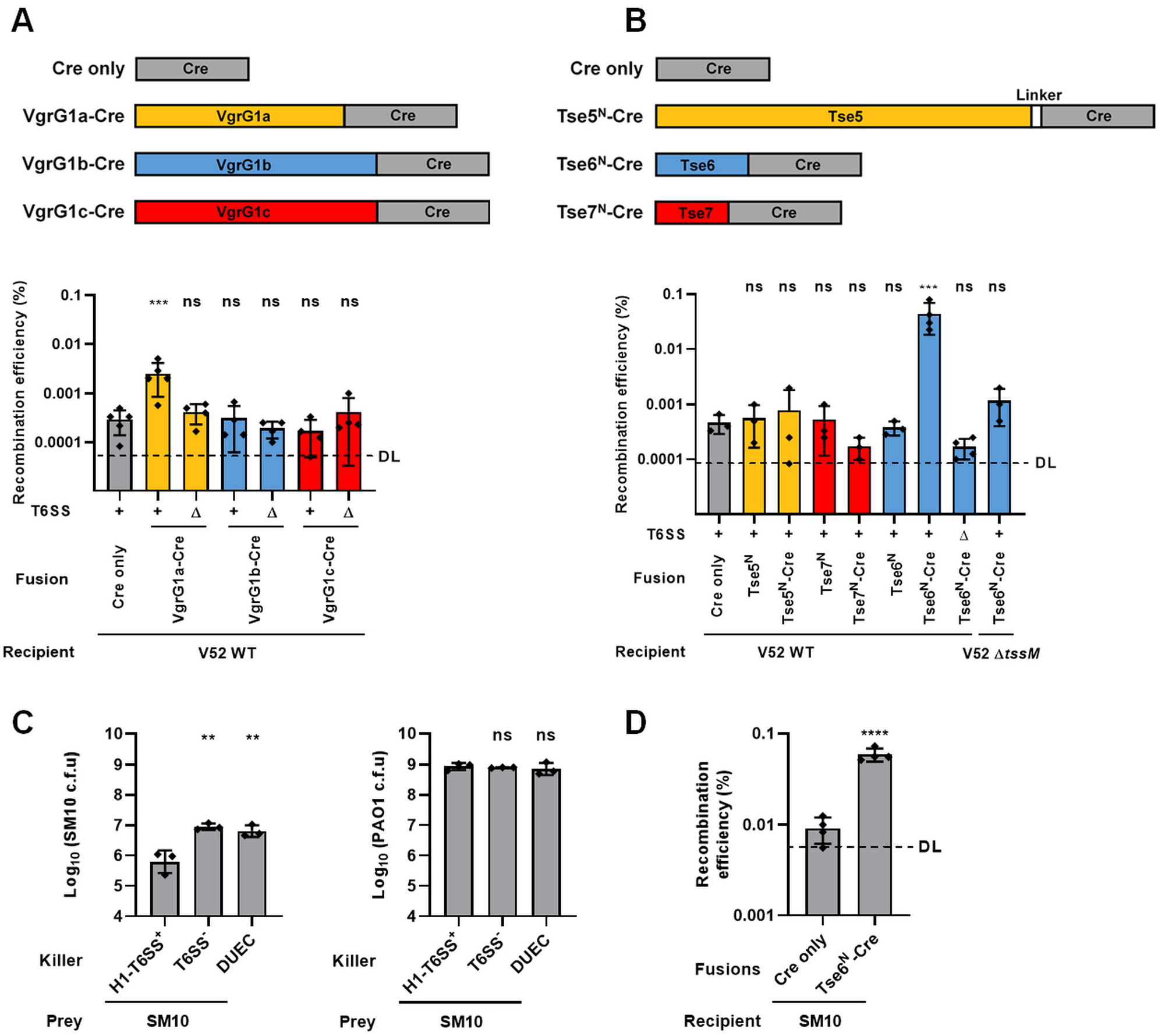
DUEC delivers a functional Cre into reporter cells. A, Diagram showing Cre recombinase (grey) fused to VgrGs secreted by H1-T6SS of *P. aeruginosa* (upper). Cre recombinase was inserted before stop codons of VgrG1a (yellow), VgrG1b (blue) and VgrG1c (red). Recombination efficiency of VgrG-Cre fusions delivered by *P. aeruginosa* eight-effector-inactivated mutant DUEC (+) or T6SS^−^ (Δ) carrying VgrG-Cre fusions after co-incubation with *V. cholerae* V52 wild type (WT) carrying pFIGR plasmid (lower). Recovery of *V. cholerae* Cre-recombined recipient cells (Gen^R^), *V. cholerae* recipient cells with pFIGR plasmid (Carb^R^), and *P. aeruginosa* mutant carrying VgrG-Cre fusions (Donor) are shown in Fig. S1A-S1C respectively. B, Diagram showing Cre recombinase (grey) fused to VgrG-dependent Tse effectors secreted by H1-T6SS of *P. aeruginosa* (upper). Cre recombinase was inserted after codon 1145 of Tse5 (yellow), codon 281 of Tse6 (blue) and codon 225 of Tse7 (red). Recombination efficiency of Tse-Cre fusions delivered by *P. aeruginosa* mutant DUEC (+) or T6SS^−^ (Δ) carrying Tse-Cre fusions after co-incubation with *V. cholerae* V52 wild type (WT) or T6SS-null (Δ*tssM*) carrying pFIGR plasmid (lower). Recovery of *V. cholerae* Cre-recombined recipient cells (Gen^R^), *V. cholerae* recipient cells with pFIGR plasmid (Carb^R^), and *P. aeruginosa* DUEC carrying Tse-Cre fusions (Donor) is shown in Fig. S2A-S2C respectively. C, Survival of *E. coli* SM10 λpir with Strp^R^ (left) and *P. aeruginosa* PAO1 mutants (right) after competition assay. Killer and prey were mixed at the ratio of 10: 1. D, Recombination efficiency of Tse6^N^-Cre fusions delivered by *P. aeruginosa* eight-effector-inactivated mutant DUEC carrying Tse6^N^-Cre fusions after co-incubation with *E. coli* SM10 λpir carrying pFIGR plasmid. Recovery of SM10 λpir Cre-recombined recipient cells (Gen^R^), SM10 λpir recipient cells with pFIGR plasmid (Carb^R^), and *P. aeruginosa* mutants carrying Cre fusions (Donor) is shown in Fig. S3A-S3C respectively. DL, approximate detection limit. Error bars indicate the mean ± standard deviation of at least three different biological duplicates. One-way ANOVA with Dunnett’s multiple comparisons test compared to H1-T6SS^+^ in C or DUEC with Cre only in A, B, and D. \*\**P* < 0.01, \*\*\**P* < 0.001, \*\*\*\**P* < 0.0001; ns, not significant. Source data are provided as a Source Data file.

### Cre delivery can be improved by fusion with the effector Tse6

Having demonstrated that Cre can be delivered by the H1-T6SS as a VgrG1a fused cargo, we next tested whether effector-fusions can also be used for Cre delivery. Because the cargo size of Cre (343 amino acids) might prevent it from getting inside the inner lumen of the Hcp tube, we only tested the three VgrG-dependent effectors (Tse5-7) as carriers. To abolish toxicity, we removed the C-terminal toxin domains of these effectors and replaced with Cre instead (Fig. 2B). We performed the delivery assay by co-incubating donor and recipient cells expressing the Tse-Cre plasmids, the Cre-only or effector-only plasmids, and the pFIGR reporter plasmid, respectively (Fig. 2B and Fig. S2A-S2C). We also included the *V. cholerae tssM* deletion mutant as a recipient control which cannot induce the dueling effect of H1-T6SS (Fig. 2B and Fig. S2A-S2C). We found that the Tse6^N^-Cre construct produced the most Gen^R^ colonies, with 100-fold higher recombination efficiency than the Tse6^N^-only or Cre-only plasmids (Fig. 2B and Fig. S2A-S2C). There were some background levels of spontaneous Gen^R^ colonies in Tse5^N^-Cre and Tse7^N^-Cre samples that are comparable to the negative control samples. Notably, neither the *tssB1* deletion mutant as donor nor the *V. cholerae tssM* deletion mutant as recipient induced Tse6^N^-Cre-mediated recombination above background levels (Fig. 2B and Fig. S2A-S2C). These results suggest that Tse6^N^ can serve as an effective secretion tag of H1-T6SS for the delivery of cargo proteins.

### DUEC delivers Tse6-Cre into conjugative *E. coli*

Next, we tested whether DUEC could deliver cargos to other species. Because RP4 conjugation can induce the assembly of H1-T6SS ^1^, we tested whether DUEC can deliver Cre into *E. coli* strain SM10 λpir Strp^R^, to which we transformed the pFIGR reporter. Using a competition assay, we first confirmed that the parental H1-T6SS^+^ cell (Δ*retS*, Δ*tssB2*, Δ*tssB3*, TssB1-sfGFP) could effectively kill *E. coli* but the DUEC strain exhibited abolished killing similar to the T6SS null mutant (Δ*retS*, Δ*tssB1*, Δ*tssB2*, Δ*tssB3*) (Fig. 2C). Then, we tested the Cre-delivery efficiency using donor strains expressing Tse6^N^-Cre or Cre-only vectors. By mixing donor and recipient cells, we found that Tse6^N^-Cre produced a 6-fold higher number of Gen^R^ recombinants than the Cre only control (Fig. 2D and Fig. S3A-S3C). These data indicate that DUEC could sense conjugation and deliver cargo into *E. coli*.

### Transmembrane domain 1 (TMD1) is essential for Tse6-Cre delivery into T6SS^+^ cells

Tse6 contains two predicted transmembrane domains TMD1 (1-61 residues) and TMD2 (180-222 residues) flanking a conserved PAAR domain. Each TMD interacts with a homodimer of EagT6 chaperone that facilitates the loading of Tse6 onto VgrG1a ^44^, likely through the interaction between PAAR and VgrG1a. To test whether the length of the Tse6^N^ secretion tag can be further reduced and whether the TMDs are essential for the Cre delivery, we deleted TMD1 and TMD2 individually (Tse6^ΔTMD1^-Cre and Tse6^ΔTMD2^-Cre), or both TMD regions (Tse6^ΔTMD1,2^-Cre) in the Tse6^N^-Cre fusion (Fig. 3A). Using the pFIGR reporter strains, the removal of TMD2 slightly reduced Tse6^N^-Cre mediated recombination while the removal of TMD1 abolished recombination, suggesting that TMD1 is required for the Tse6 delivery while the TMD2 domain is nonessential (Fig. 3B and Fig. S4A-S4C). Although the removal of TMD2 did not result in increased recombination efficiency, its reduced size might be useful for the delivery of larger cargo proteins.

**Fig. 3.**
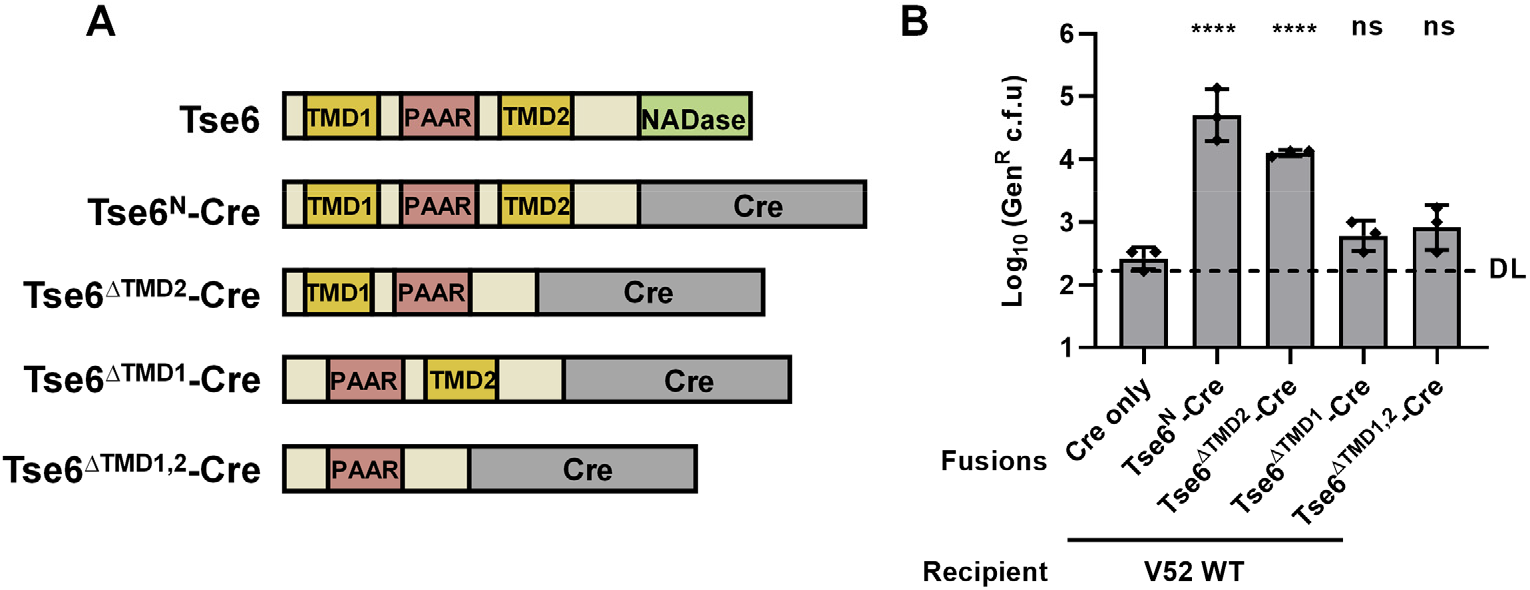
TMD1 of Tse6 is essential for Cre delivery by *P. aeruginosa*. A, Diagram showing Cre recombinase (grey) fused to different Tse6 mutants. Truncated Tse6 lacks either one TMD (Tse6^ΔTMD2^: deletion of 180-222 residues; Tse6^ΔTMD1^: deletion of 1-61 residues) or both TMD (Tse6^ΔTMD1,2^). B, Recovery of *V. cholerae* Cre-recombined recipient cells (Gen^R^) after co-incubation of *P. aeruginosa* eight-effector-inactivated DUEC mutant carrying truncated Tse6-Cre fusions with *V. cholerae* V52 wild type (WT) carrying pFIGR plasmid. Recombination efficiency, recovery of *V. cholerae* recipient cells with pFIGR plasmid (Carb^R^), and *P. aeruginosa* mutants carrying truncated Tse6-Cre fusions (Donor) is shown in Fig. S4A-S4C respectively. DL, approximate detection limit. Error bars indicate the mean ± standard deviation of at least three different biological duplicates. One-way ANOVA with Dunnett’s multiple comparisons test compared DUEC with Cre only fusion. \*\*\*\**P* < 0.0001; ns, not significant. Source data are provided as a Source Data file.

### DUEC can selectively deliver to T6SS^+^ cells in a mixed community

To test whether DUEC could selectively deliver effectors into T6SS^+^ cells within a mixed population, we co-incubated DUEC carrying Tse6^N^-Cre with both V52 and its Δ*tssM* mutant carrying the pFIGR reporter (Fig. 4A). Out of over a hundred recombinants, we randomly selected 10 colonies and tested whether these were T6SS^+^ cells using an *E. coli* prey expressing a luciferase reporter in the competition assay. We found that all tested recombinants were T6SS^+^ wild type (Fig. 4B and Fig. S5A-S5D), indicating that DUEC could differentiate T6SS^+^ cells in a mixed community.

**Fig. 4.**
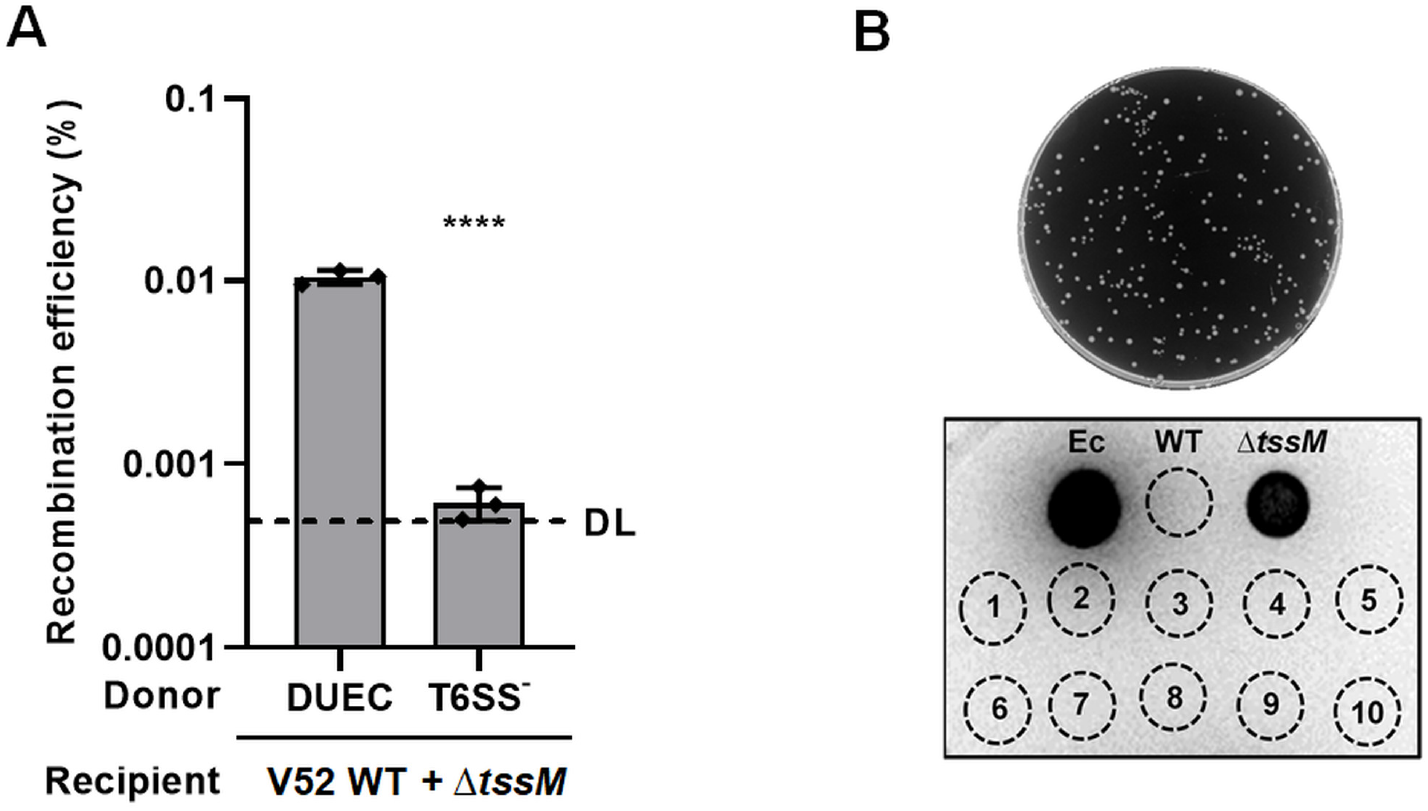
DUEC responds to T6SS^+^ cells in a mixed community. A, Recombination efficiency of Tse6^N^-Cre fusions delivered by *P. aeruginosa* DUEC or T6SS^−^ mutant carrying Tse6^N^-Cre fusions after co-incubation of *P. aeruginosa* mutants carrying Tse6^N^-Cre fusions with both *V. cholerae* V52 wild type (WT) and T6SS-null (Δ*tssM*) carrying pFIGR plasmids at the ratio of 10: 1: 1. Recovery of *V. cholerae* Cre-recombined recipient cells (Gen^R^), *V. cholerae* recipient cells with pFIGR plasmid (Carb^R^), and *P. aeruginosa* DUEC carrying Tse6^N^-Cre fusion (Donor) is shown in Fig. S5A-S5C respectively. A two-tailed Student’s *t*-test compared to DUEC with Tse6^N^-Cre was used. Error bars indicate the mean ± standard deviation of at least three different biological duplicates. Source data are provided as a Source Data file. B, Recovery of *V. cholerae* Cre-recombined recipient (Gen^R^) cells after co-incubation of *P. aeruginosa* DUEC carrying Tse6^N^-Cre with both *V. cholerae* WT and the Δ*tssM* mutant carrying the pFIGR plasmid at the ratio of 10: 1: 1 (upper). A representative luminescent image of *E. coli* MG1655 with pBAD18kan-lux plasmid after attacked by different *V. cholerae* strains (lower). Ec: *E. coli* MG1655 with pBAD18kan-lux; WT: *V. cholerae* wild type; Δ*tssM*: *V. cholerae* T6SS-null mutant; *V. cholerae* Gen^R^ isolated colonies (1 to 10) after co-incubation of *V. cholerae* WT, Δ*tssM* carrying pFIGR plasmid with *P. aeruginosa* DUEC carrying Tse6^N^-Cre fusion. Image of mixtures of *E. coli* MG1655 with pBAD18kan-lux plasmid with different *V. cholerae* strains is shown in Fig. S5D.

## Discussion

The versatility of microbial environmental sensing has provided numerous valuable tools for conditional gene expression in synthetic biology and biotechnology applications. However, it remains challenging to precisely differentiate and modulate a select member within a complex microbial community. Here, we have constructed the DUEC platform, a dueling-competent and detoxified *P. aeruginosa* H1-T6SS that can translocate proteins into the cytosol of a neighboring cell when sensing a physical disturbance signal exerted by the neighboring cell. Although effector-inactivating strategies have been recently employed ^9,19^, the T6SSs in *P. aeruginosa* are far more complex not only for having three independent clusters but also for the relatively large number of effectors with different routes of delivery ^30,33^. For cargo protein delivery, we determine a secretion tag, a truncated Tse6, that could effectively deliver the Cre recombinase by DUEC to the cytosol of T6SS^+^ *V. cholerae* cells. Collectively, our data demonstrate a functional inter/intraspecies direct-contact sensing module that can be used for targeted protein delivery or, coupled with cargo functions, to control cargo-dependent downstream modules (Fig. 5).

**Fig. 5.**
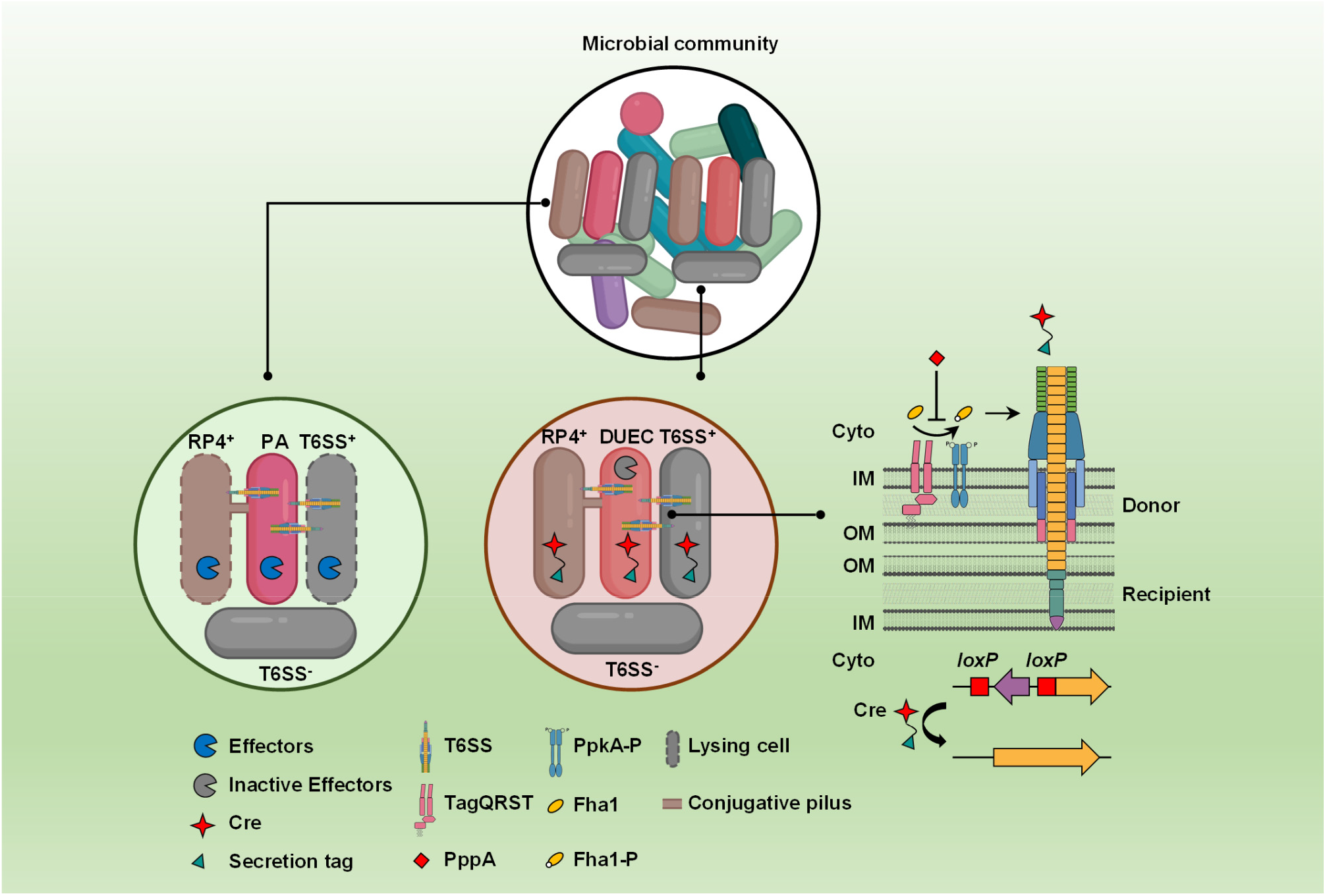
A schematic model of DUEC for precise cargo delivery within a mixed population. Left, *P.aeruginosa* wild type (PA) assembles its H1-T6SS after sensing the attack from neighboring T6SS^+^ cells or RP4 conjugation, and delivers toxic effectors into the attackers, resulting in cell lysis (dotted lysing cell). Middle, *P. aeruginosa* DUEC with active T6SS but inactivated effectors does not kill T6SS^+^ cells or conjugative *E. coli* cells, but can precisely respond to T6SS^+^ cells in a mixed community. Right, the dueling/tit-for-tat signal transduction module comprises the signal-sensing of TagQRST and the subsequent phosphorylation of Fha1 by the PpkA kinase, which leads to T6SS assembly in *P. aeruginosa* DUEC and the resultant delivery of Cre into recipient cells. The secretion-tag fused Cre remains active and mediates specific recombination. Thus, the post-translational dueling/tit-for-tat sensing module is now equipped with a powerful genetic switch for broad downstream outputs.

The dueling response module offers probably one of the fastest signal-transduction modules for it can respond to a number of membrane-disturbing signals, including EDTA and polymyxin B, within seconds ^1,42^. However, it is controlled by a cascade of signal-sensing and phosphorelay proteins with a direct outcome of T6SS assembly, lacking a DNA-binding regulator that is commonly used in synthetic biology modules. The complexity of this signal relay and output presents a considerable challenge for heterologous expression and adoption of this cascade for controlled gene expression. Combining effector-inactivation to preserve the H1-T6SS activity and Cre delivery, we demonstrate a prototypical regulatory module that can sense a physical puncture trigger a genetic change, in this case, gain of antibiotic resistance through Cre-mediated recombination. Both the cargo Cre and the combination outcome can be flexibly replaced with other enzymes or output modules, offering an expanded potential for broad application of the DUEC system.

## Materials and Methods

### Bacterial strains and growth conditions

Strains, plasmids, and primers used in this study are listed in Table S1. Cultures were grown in LB (1% [w/v] tryptone, 0.5% [w/v] yeast extract, 0.5% [w/v] NaCl) aerobically at 37 °C. Antibiotics were used at the following concentrations: streptomycin (100 μg/ml), irgasan (25 μg/ml), gentamicin (20 μg/ml), carbenicillin (100 μg/ml) and kanamycin (50 μg/ml).

Plasmids were constructed using Gibson assembly ^45^ and overlapping PCR. All constructs were confirmed by Sanger sequencing.

### Construction of mutants

All chromosomal mutants were constructed by crossover PCR and homologous recombination using suicide vector pEXG2.0 as previously described ^5^. Homologous arms were amplified and overlapped by PCR and cloned into pEXG2.0 by Gibson assembly. The suicide vectors carrying homologous arms were first transformed into donor *E. coli* WM6026 or SM10 λpir by transformation, and then to the recipient strains by conjugation. Trans-conjugants were selected on LB plates with antibiotics and sucrose-resistant colonies were selected on 6% sucrose plates. In-frame deletions were confirmed by PCR and site-specific mutagenesis was confirmed by sequencing.

### Bacterial cell killing assay

*P. aeruginosa* cells were grown overnight on LB-agar plates with appropriate antibiotics. The killer cells were normalized to OD_600_ =1 and incubated at 37 °C for 1 h, 1/50 (for *P. aeruginosa*) sub-cultured to OD_600_ =1. Overnight cultures of *V. cholerae* or *E. coli* SM10 λpir cells were normalized to OD_600_ =1. Cells were centrifuged at 10, 000 × *g* for 30 seconds twice, resuspended in fresh LB, mixed at a ratio of 5: 1 (killer: prey), and spotted on LB-agar plates. After co-incubation for 3 h at 37 °C, the bacteria were resuspended in 500 μl fresh LB and a series of 10-fold serial dilutions were performed and plated on LB-agar with the appropriate antibiotic to assess the survival of killer and prey cells. LB-agar with 100 μg/ml streptomycin to select for *V. cholerae* V52 or *E. coli* SM10 λpir, and 25 μg/ml irgasan for recovery of *P. aeruginosa*. All killing assays were done at least in triplicate. The mean Log_10_ c.f.u of recovered prey were plotted, and error bars show the mean ± standard deviation of three biological replicates. One-way ANOVA with Dunnett’s multiple comparisons test was used to determine *P*-values.

### Screening of SM10 λpir mutants with spontaneous resistance to streptomycin

*E. coli* SM10 λpir cells were cultured for about 12 h at 37 °C to stationary phase and 2 ml cells were collected at 10, 000 × *g* for 30 seconds twice, and streaked on LB plates with 100 μg/ml streptomycin. Single colonies were picked after overnight culture at 37 °C.

### Cre delivery assay

*P. aeruginosa* cells were grown overnight on LB-agar plates with appropriate antibiotics, 1/50 sub-cultured and incubated at 37 °C for ~3 h to OD_600_ =1. *P. aeruginosa* cells were centrifuged at 10, 000 × *g* for 30 seconds twice, resuspended in fresh LB to OD_600_ =10. Overnight cultures of prey cells (*V. cholerae* strains or *E. coli* SM10 λpir strain with pFIGR) were normalized to OD_600_ =1. *P. aeruginosa* and *V. cholerae* or *E. coli* SM10 λpir cells were mixed at a ratio of 10: 1 (donor: recipient) and spotted on LB-agar plates with 1 mM isopropyl-β-D-thiogalactoside (IPTG) to induce the expression of Cre fusions. After co-incubation for 3 h at 37 °C, the bacteria were resuspended in 500 μl fresh LB and a series of 10-fold serial dilutions were performed and plated on LB-agar with the appropriate antibiotic to assess the survival of killer and prey cells. LB-agar with 100 μg/ml streptomycin and 20 μg/ml gentamicin were used to select for *V. cholerae* or *E. coli* SM10 λpir with obtained gentamicin resistance, 100 μg/ml streptomycin and 100 μg/ml carbenicillin for original *V. cholerae* or *E. coli* SM10 λpir recipient cells and 25 μg/ml irgasan and 20 μg/ml gentamicin for recovery of *P. aeruginosa*. For Cre delivery in a mixed community, *P. aeruginosa* and *V. cholerae* cells were mixed at a ratio of 10:1:1 (donor: recipient WT: recipient Δ*tssM*). 10 random colonies were picked from *V. cholerae* cells with obtained gentamicin resistance for competition assay of *V. cholerae* Gen^R^ cells against *E. coli* MG1655 with a luciferase reporter. Data are shown as either Log_10_ c.f.u recovered on LB plates with appropriate antibiotics or as recombination efficiency, calculated as Gen^R^ c.f.u/Carb^R^ c.f.u. If no colonies were observed, samples were listed as the detection limit (DL), shown as if 0.5 colonies were counted ^22^. Error bars show the mean ± standard deviation of three biological replicates. One-way ANOVA with Dunnett’s multiple comparisons test was used to determine *P*-values.

### Fluorescence microscopy image acquisition and analysis

Single colonies were grown overnight and diluted 1/50 in LB with 25 μg/ml irgasan for *P. aeruginosa* mutants and cells were grown to OD_600_~0.8-1. *P. aeruginosa* cells were centrifuged at 10, 000 × *g* for 30 seconds twice, resuspended in fresh LB to OD_600_ =10. Cells were spotted on 1% agarose-0.5 × PBS pads. Regions on the edges were taken. The microscopy images were obtained using a Nikon Ti-E inverted microscope with a Perfect Focus System (PFS)and a CFI Plan Apochromat Lambda 100 × oil objective lens. ET-GFP (Chroma 49002) filter sets were used for the GFP fluorescence signal.

All images were analyzed and manipulated using Fiji software. Images from a 5-min series were normalized to the same mean intensity to correct for photobleaching, as described previously ^46^. The plugin “Temporal-Color code” for Fiji was used to assess the active sheath over the time frame. All imaging experiments were performed with at least three biological replicates.

### Software

Statistical analysis was treated using the GraphPad Prism software (8.2.1).

## Data availability

The source data underlying Figs. 1C, 2A-2D, 3B, 4A and Figs. S1A-S1C, S2A-S2C, S3A-S3C, S4A-S4C and S5A-S5C are provided as a Source Data file. Other data supporting the findings of this study are available within the paper or from the corresponding author upon request.

## Acknowledgments

This work was supported by funding from the National Key R&D Program of China (2020YFA0907200), National Natural Science Foundation of China (32030001), and National Natural Science Foundation of China-Youth Science Fund (82102399). We thank Kevin Manera and Steve Hersch for their helpful comments. The funders had no role in study design, data collection and interpretation, or the decision to publish.

## Author contributions

T.D. conceived the project. L.W. performed most experiments, analyzed results and prepared all Figures. L.W., M.T., T.P., and S.Y. contributed to competition assay. H.L. contributed to strain construction. X.L. contributed to the experimental design. L.W., S.S., X.L., and T.D. wrote the paper.

## Conflict of Interest

The authors declare no conflict of interest.

**Correspondence and request for materials should be addressed to T. Dong.**

## References

1. Ho, B. T., Basler, M. & Mekalanos, J. J. Type 6 secretion system-mediated immunity to type 4 secretion system-mediated gene transfer. Science 342, 250–253 (2013).

2. Basler, M. & Mekalanos, J. J. Type 6 secretion dynamics within and between bacterial cells. Science 337, 815 (2012).

3. Basler, M., Ho, B. T. & Mekalanos, J. J. Tit-for-tat: type VI secretion system counterattack during bacterial cell-cell interactions. Cell 152, 884–894 (2013).

4. Pukatzki, S. et al. Identification of a conserved bacterial protein secretion system in *Vibrio cholerae* using the *Dictyostelium* host model system. Proc. Natl. Acad. Sci. U S A 103, 1528–1533 (2006).

5. Mougous, J. D. et al. A virulence locus of *Pseudomonas aeruginosa* encodes a protein secretion apparatus. Science 312, 1526–1530 (2006).

6. Ho, B. T., Dong, T. G. & Mekalanos, J. J. A view to a kill: the bacterial type VI secretion system. Cell Host Microbe 15, 9–21 (2014).

7. Vettiger, A. & Basler, M. Type VI secretion system substrates are transferred and reused among sister cells. Cell 167, 99–110 (2016).

8. Ho, B. T., Fu, Y., Dong, T. G. & Mekalanos, J. J. *Vibrio cholerae* type 6 secretion system effector trafficking in target bacterial cells. Proc. Natl. Acad. Sci. U S A 114, 9427–9432 (2017).

9. Liang, X. et al. An onboard checking mechanism ensures effector delivery of the type VI secretion system in *Vibrio cholerae*. Proc. Natl. Acad. Sci. U S A 116, 23292–23298 (2019).

10. Pei, T. T. et al. Delivery of an Rhs-family nuclease effector reveals direct penetration of the gram-positive cell envelope by a type VI secretion system in *Acidovorax citrulli*. mLife 1, 66–78 (2022).

11. Pei, T. T. et al. Intramolecular chaperone-mediated secretion of an Rhs effector toxin by a type VI secretion system. Nat. Commun. 11, 1865 (2020).

12. Trunk, K. et al. The type VI secretion system deploys antifungal effectors against microbial competitors. Nat. Microbiol. 3, 920–931 (2018).

13. Hood, R. D. et al. A type VI secretion system of *Pseudomonas aeruginosa* targets a toxin to bacteria. Cell Host Microbe 7, 25–37 (2010).

14. MacIntyre, D. L., Miyata, S. T., Kitaoka, M. & Pukatzki, S. The *Vibrio cholerae* type VI secretion system displays antimicrobial properties. Proc. Natl. Acad. Sci. U S A 107, 19520–19524 (2010).

15. Si, M. et al. Manganese scavenging and oxidative stress response mediated by type VI secretion system in *Burkholderia thailandensis*. Proc. Natl. Acad. Sci. U S A 114, E2233–E2242 (2017).

16. Song, L. et al. Contact-independent killing mediated by a T6SS effector with intrinsic cell-entry properties. Nat. Commun. 12, 423 (2021).

17. Toska, J., Ho, B. T. & Mekalanos, J. J. Exopolysaccharide protects *Vibrio cholerae* from exogenous attacks by the type 6 secretion system. Proc. Natl. Acad. Sci. U S A 115, 7997–8002 (2018).

18. Wu, C. et al. Effector loading onto the VgrG carrier activates type VI secretion system assembly. EMBO Rep. 21, e47961 (2020).

19. Liang, X. et al. Characterization of lysozyme-like effector TseP reveals the dependence of type VI secretion system (T6SS) secretion on effectors in *Aeromonas dhakensis* strain SSU. Appl. Environ. Microbiol. 87, e00435–21 (2021).

20. Liang, X. et al. VgrG-dependent effectors and chaperones modulate the assembly of the type VI secretion system. PLoS Pathog. 17, e1010116 (2021).

21. Ma, A. T., McAuley, S., Pukatzki, S. & Mekalanos, J. J. Translocation of a *Vibrio cholerae* type VI secretion effector requires bacterial endocytosis by host cells. Cell Host Microbe 5, 234–243 (2009).

22. Hersch, S. J., Lam, L. & Dong, T. G. Engineered type six secretion systems deliver active exogenous effectors and Cre recombinase. mBio 12, e0111521 (2021).

23. Sana, T. G., Berni, B. & Bleves, S. The T6SSs of *Pseudomonas aeruginosa* strain PAO1 and their effectors: beyond bacterial-cell targeting. Front. Cell. Infect. Microbiol. 6, 61 (2016).

24. Sana, T. G. et al. The second type VI secretion system of *Pseudomonas aeruginosa* strain PAO1 is regulated by quorum sensing and Fur and modulates internalization in epithelial cells. J. Biol. Chem. 287, 27095–27105 (2012).

25. Wettstadt, S., Wood, T. E., Fecht, S. & Filloux, A. Delivery of the *Pseudomonas aeruginosa* phospholipase effectors PldA and PldB in a VgrG- and H2-T6SS-dependent manner. Front. Microbiol. 10, 1718 (2019).

26. Burkinshaw, B. J. et al. A type VI secretion system effector delivery mechanism dependent on PAAR and a chaperone-co-chaperone complex. Nat. Microbiol. 3, 632–640 (2018).

27. Lin, J. et al. A *Pseudomonas* T6SS effector recruits PQS-containing outer membrane vesicles for iron acquisition. Nat. Commun. 8, 14888 (2017).

28. Howard, S. A. et al. The breadth and molecular basis of Hcp-driven type VI secretion system effector delivery. mBio 12, e00262–21 (2021).

29. Russell, A. B. et al. Type VI secretion delivers bacteriolytic effectors to target cells. Nature 475, 343–349 (2011).

30. LaCourse, K. D. et al. Conditional toxicity and synergy drive diversity among antibacterial effectors. Nat. Microbiol. 3, 440–446 (2018).

31. Whitney, J. C. et al. Genetically distinct pathways guide effector export through the type VI secretion system. Mol. Microbiol. 92, 529–542 (2014).

32. Whitney, J. C. et al. An interbacterial NAD(P)^+^ glycohydrolase toxin requires elongation factor Tu for delivery to target cells. Cell 163, 607–619 (2015).

33. Nolan, L. M. et al. Identification of Tse8 as a type VI secretion system toxin from *Pseudomonas aeruginosa* that targets the bacterial transamidosome to inhibit protein synthesis in prey cells. Nat. Microbiol. 6, 1199–1210 (2021).

34. Pissaridou, P. et al. The *Pseudomonas aeruginosa* T6SS-VgrG1b spike is topped by a PAAR protein eliciting DNA damage to bacterial competitors. Proc. Natl. Acad. Sci. U S A 115, 12519–12524 (2018).

35. Pedrolli, D. B. et al. Engineering microbial living therapeutics: the synthetic biology toolbox. Trends Biotechnol. 37, 100–115 (2019).

36. Cubillos-Ruiz, A. et al. Engineering living therapeutics with synthetic biology. Nat. Rev. Drug Discov. 20, 941–960 (2021).

37. Bartley, B. A., Kim, K., Medley, J. K. & Sauro, H. M. Synthetic biology: engineering living systems from biophysical principles. Biophys. J. 112, 1050–1058 (2017).

38. Khalil, A. S. & Collins, J. J. Synthetic biology: applications come of age. Nat. Rev. Genet. 11, 367–379 (2010).

39. Mougous, J. D., Gifford, C. A., Ramsdell, T. L. & Mekalanos, J. J. Threonine phosphorylation post-translationally regulates protein secretion in *Pseudomonas aeruginosa*. Nat. Cell Biol. 9, 797–803 (2007).

40. Hsu, F., Schwarz, S. & Mougous, J. D. TagR promotes PpkA-catalysed type VI secretion activation in *Pseudomonas aeruginosa*. Mol. Microbiol. 72, 1111–1125 (2009).

41. Casabona, M. G. et al. An ABC transporter and an outer membrane lipoprotein participate in posttranslational activation of type VI secretion in *Pseudomonas aeruginosa*. Environ. Microbiol. 15, 471–486 (2013).

42. Wilton, M. et al. Chelation of membrane-bound cations by extracellular DNA activates the type VI secretion system in *Pseudomonas aeruginosa*. Infect. Immun. 84, 2355–2361 (2016).

43. Pukatzki, S., Ma, A. T., Revel, A. T., Sturtevant, D. & Mekalanos, J. J. Type VI secretion system translocates a phage tail spike-like protein into target cells where it cross-links actin. Proc. Natl. Acad. Sci. U S A 104, 15508–15513 (2007).

44. Quentin, D. et al. Mechanism of loading and translocation of type VI secretion system effector Tse6. Nat. Microbiol. 3, 1142–1152 (2018).

45. Gibson, D. G. et al. Enzymatic assembly of DNA molecules up to several hundred kilobases. Nat. Methods 6, 343–345 (2009).

46. Wong, M. J. Q. et al. Microbial herd protection mediated by antagonistic interaction in polymicrobial communities. Appl. Environ. Microbiol. 82, 6881–6888 (2016).

