## Supplemental materials for "A dueling-competent signal-sensing module guides precise delivery of cargo proteins into target cells by engineered *Pseudomonas aeruginosa*"

Supplementary Figure 1:

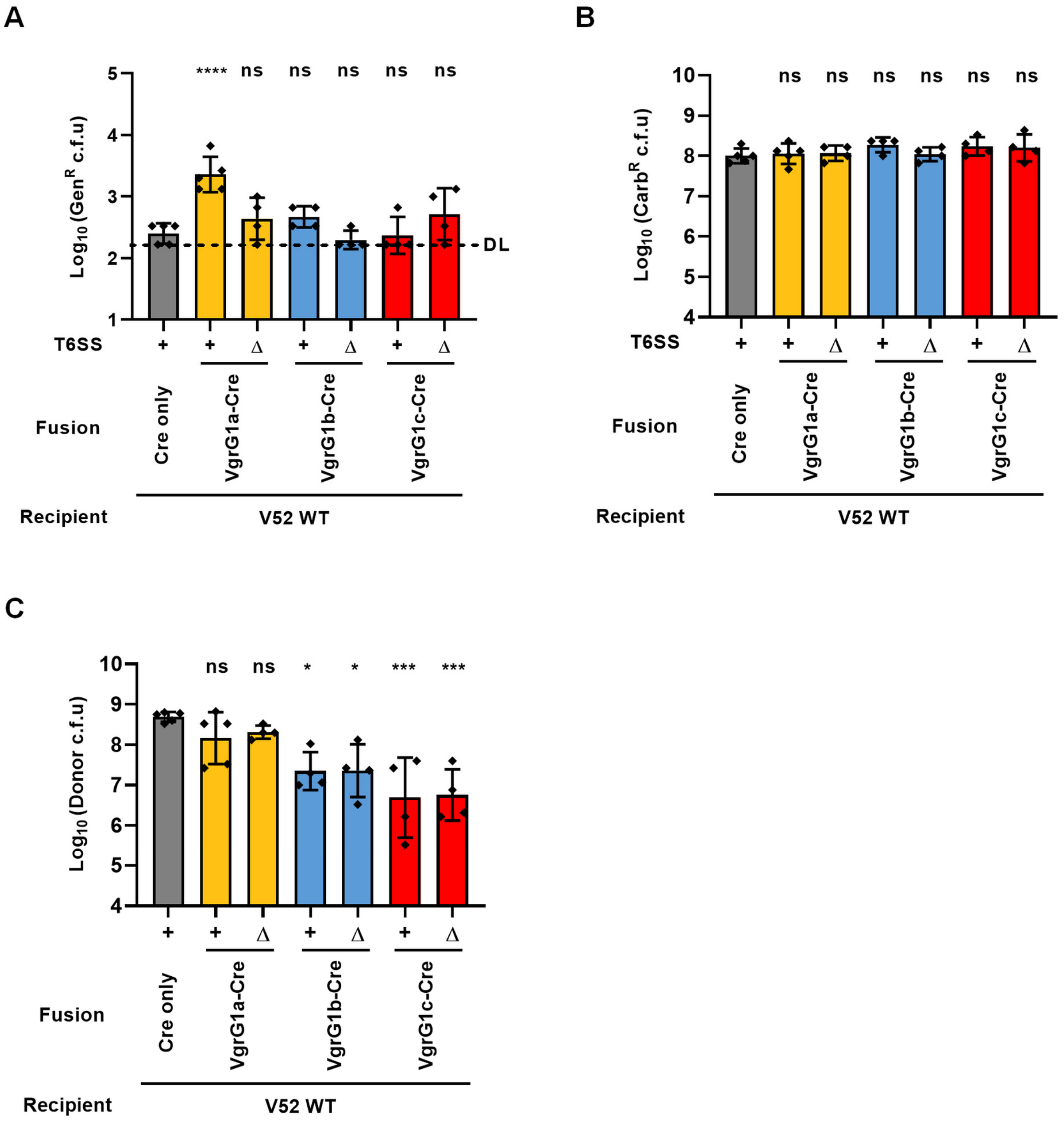


**Supplementary Figure 1. VgrG1a-Cre fusions can be delivered into recipient cells in a T6SS-dependent manner.** A to C, Recovery of *V. cholerae* Cre-recombined recipient cells (Gen$^{R}$, A), *V. cholerae* recipient cells with pFIGR plasmid (Carb$^{R}$, B), and *P. aeruginosa* mutants carrying VgrG-Cre fusions (Donor, C) after co-incubation of *P. aeruginosa* mutants carrying VgrG-Cre fusions with *V. cholerae* wild type (WT) carrying pFIGR plasmid. Recombination efficiency of VgrG-Cre fusions is shown in Fig. 2A. DL, approximate detection limit. Error bars indicate the mean ± standard deviation of at least three different biological duplicates. One-way ANOVA Dunnett's multiple-comparison test compared DUEC with Cre only. *$P < 0.05$, ***$P < 0.001$, ****$P < 0.0001$; ns, not significant. Source data are provided as a Source Data file.

Supplementary Figure 2:

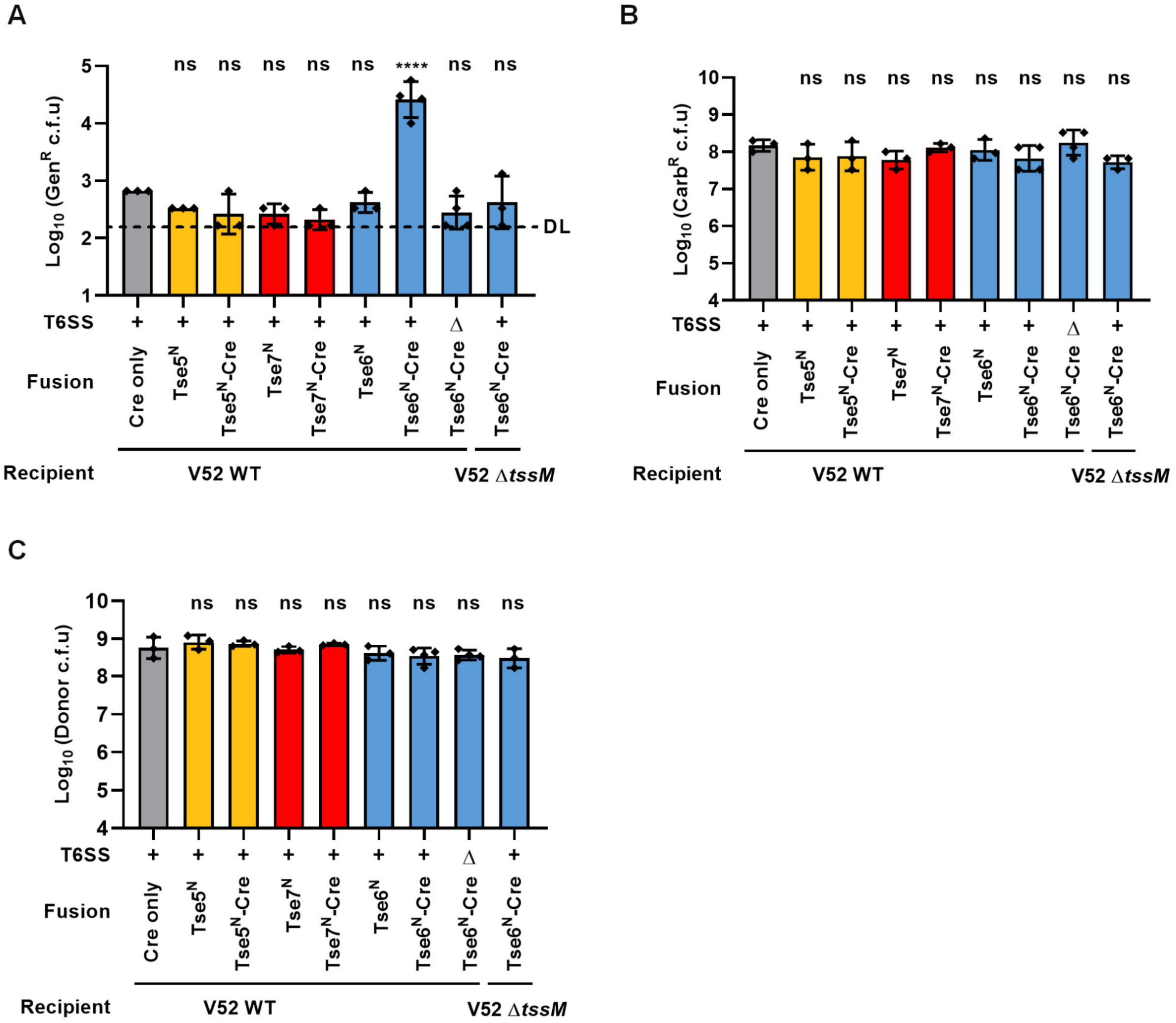


**Supplementary Figure 2. Active Cre can be delivered as effector fusions by *P. aeruginosa* DUEC.** A to C, Recovery of *V. cholerae* Cre-recombined recipient cells ($Gen^R$, A), *V. cholerae* recipient cells with pFIGR plasmid ($Carb^R$, B), and *P. aeruginosa* mutant carrying Tse-Cre fusions (Donor, C) after co-incubation of *P. aeruginosa* eight-effector-inactivated mutant DUEC carrying Tse-Cre fusions with *V. cholerae* V52 wild type (WT) or T6SS-null (Δ*tssM*) carrying pFIGR plasmid. Recombination efficiency of Tse-Cre fusions is shown in Fig. 2B. DL, approximate detection limit. Error bars indicate the mean ± standard deviation of at least three different biological duplicates. One-way ANOVA with Dunnett's multiple comparisons test compared DUEC with Cre only. ****$P < 0.0001$; ns, not significant. Source data are provided as a Source Data file.

Supplementary Figure 3

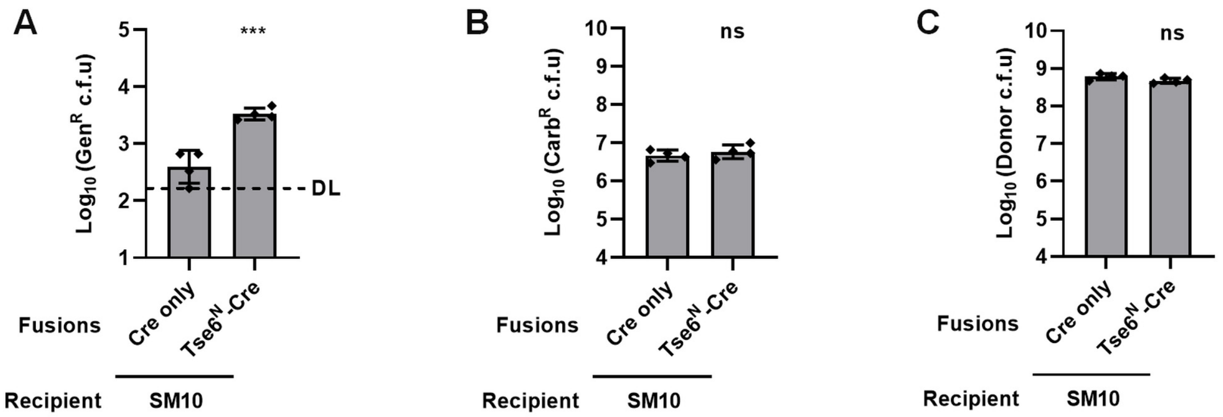


**Supplementary Figure 3. DUEC delivers Tse6-Cre into *E. coli*.** A to C, Recovery of *E. coli* SM10 Cre-recombined recipient cells ($Gen^R$, A), *E. coli* SM10 recipient cells with pFIGR plasmid ($Carb^R$, B), and *P. aeruginosa* DUEC $Tse6^N$-Cre fusions (Donor, C) after co-incubation of *P. aeruginosa* DUEC mutant carrying $Tse6^N$-Cre fusions with *E. coli* SM10 carrying pFIGR plasmid. Recombination efficiency is shown is Fig. 2D. DL, approximate detection limit. Error bars indicate the mean ± standard deviation of at least three different biological duplicates. ***$P < 0.001$; ns, not significant. Source data are provided as a Source Data file.

Supplementary Figure 4

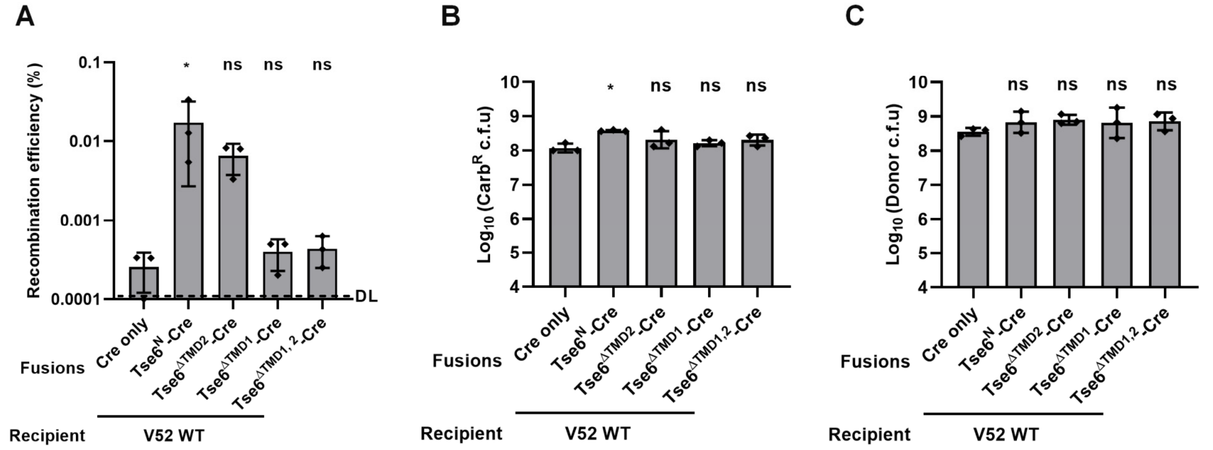


**Supplementary Figure 4. TMD1 of Tse6 is essential for Cre delivery.** A to C, Recombination efficiency (A), recovery of *V. cholerae* recipient cells with pFIGR plasmid (Carb$^R$, B), and *P. aeruginosa* DUEC carrying truncated Tse6-Cre fusions (Donor, C) after co-incubation of *P. aeruginosa* DUEC mutant carrying truncated Tse6-Cre fusions with *V. cholerae* V52 wild type (WT) carrying pFIGR plasmid. Recovery of *V. cholerae* Cre-recombined recipient cells (Gen$^R$) is shown in Fig. 3B. DL, approximate detection limit. Error bars indicate the mean ± standard deviation of at least three different biological duplicates. One-way ANOVA with Dunnett's multiple comparisons test compared to DUEC with Cre. *$P < 0.05$; ns, not significant. Source data are provided as a Source Data file.

Supplementary Figure 5

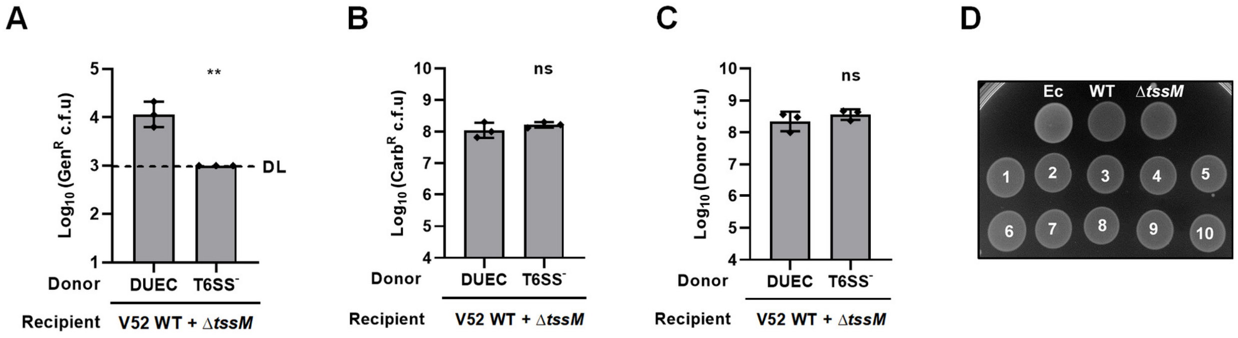


**Supplementary Figure 5. DUEC responds to T6SS$^+$ cells in a mixed community.** A to C, Recovery of *V. cholerae* Cre-recombined recipient cells (Gen$^R$, A), *V. cholerae* recipient cells with pFIGR plasmid (Carb$^R$, B), and *P. aeruginosa* DUEC or T6SS$^-$ mutant carrying Tse6$^N$-Cre fusion (Donor, C) after co-incubation of *P. aeruginosa* mutants carrying Tse6$^N$-Cre with both *V. cholerae* WT and the Δ*tssM* mutant carrying the pFIGR plasmid. Recombination efficiency of Tse6$^N$-Cre fusions is shown in Fig. 4A. D, Image of mixtures of *E. coli* MG1655 with pBAD18kan-lux plasmid with different *V. cholerae* strains. Ec: *E. coli* MG1655 with pBAD18kan-lux; WT: *V. cholerae* wild type; Δ*tssM*: *V. cholerae* T6SS-null mutant; *V. cholerae* Gen$^R$ isolated colonies (1 to 10). Luminescent image is shown in Fig. 4B. DL, approximate detection limit. Error bars indicate the mean ± standard deviation of at least three different biological duplicates. A two-tailed Student's *t*-test compared to DUEC with Tse6$^N$-Cre fusion was used in A to C. $^{**}P < 0.01$; ns, not significant. Source data are provided as a Source Data file.

**Table S1. Plasmid, strains and primers**

| Plasmid | Description | Reference |
|---|---|---|
| pPSV37-Cre | IPTG inducible expression of Cre with a C-terminal FLAG tag | This study |
| pPSV37-VgrG1a | IPTG inducible expression of full-length VgrG1a with a C-terminal FLAG tag | This study |
| pPSV37-VgrG1a-Cre | IPTG inducible expression of VgrG1a-Cre with a C-terminal FLAG tag | This study |
| pPSV37-VgrG1b | IPTG inducible expression of full-length VgrG1b with a C-terminal FLAG tag | This study |
| pPSV37-VgrG1b-Cre | IPTG inducible expression of VgrG1b-Cre with a C-terminal FLAG tag | This study |
| pPSV37-VgrG1c | IPTG inducible expression of full-length VgrG1c with a C-terminal FLAG tag | This study |
| pPSV37-VgrG1c-Cre | IPTG inducible expression of VgrG1c-Cre with a C-terminal FLAG tag | This study |
| pPSV37-Tse5$^{N}$ | IPTG inducible expression of Tse5$^{1\text{-}1145aa}$ with a C-terminal FLAG tag | This study |
| pPSV37-Tse5$^{N}$-Cre | IPTG inducible expression of Tse5$^{1\text{-}1145aa}$-Cre with a C-terminal FLAG tag | This study |
| pPSV37-Tse6$^{N}$ | IPTG inducible expression of Tse6$^{1\text{-}281aa}$ with a C-terminal FLAG tag | This study |
| pPSV37-Tse6$^{N}$-Cre | IPTG inducible expression of Tse6$^{1\text{-}281aa}$-Cre with a C-terminal FLAG tag | This study |
| pPSV37-Tse7$^{N}$ | IPTG inducible expression of Tse7$^{1\text{-}225aa}$ with a C-terminal FLAG tag | This study |
| pPSV37-Tse7$^{N}$-Cre | IPTG inducible expression of Tse7$^{1\text{-}225aa}$-Cre with a C-terminal FLAG tag | This study |
| pPSV37-Tse6$^{\Delta TMD2}$-Cre | IPTG inducible expression of Tse6$^{1\text{-}281aa}$-Cre with the deletion of TMD2 (180-222aa) and a C-terminal FLAG tag | This study |
| pPSV37-Tse6$^{\Delta TMD1}$-Cre | IPTG inducible expression of Tse6$^{1\text{-}281aa}$-Cre with the deletion of TMD1 (1-61aa) and a C-terminal FLAG tag | This study |
| pPSV37-Tse6$^{\Delta TMD1,2}$-Cre | IPTG inducible expression of Tse6$^{1\text{-}281aa}$-Cre with the deletion of TMD1 (1-61aa) and TMD2 (180-222aa) and a C-terminal FLAG tag | This study |
| pFIGR | Plasmid contains a floxed (flanked by *loxP* sites) ampicillin resistance (Carb$^{R}$) cassette that interrupts translation of a gentamicin resistance (Gent$^{R}$) gene | [1] |
| pBAD18kan-lux | Plasmid contains *luxCDABE* which can constitutively express | This study |

| Strain | Genotype | Description | Reference |
|---|---|---|---|
| *Pseudomonas aeruginosa* PAO1 | Δ*retS* Δ*tssB2* Δ*tssB3* TssB1-sfGFP | Parental strain, short for H1-T6SS$^{+}$ | This study |
| | Δ*retS* Δ*tssB2* Δ*tssB3* TssB1-sfGFP *tse1*$^{C30A}$ *tse2*$^{VK\text{-}AA}$ *tse3*$^{E250Q}$ *tse4*$^{G176V}$ Δ*tse5* *tse6*$^{D396A}$ *tse7*$^{HH\text{-}AA}$ *tse8*$^{S186A}$ | H1-T6SS$^{+}$ with chromosomal mutations of the Tse1 catalytic residue C30 to alanine, Tse2 catalytic residues V109 and K110 to alanine, Tse3 catalytic residue E250 to glutamine, Tse4 catalytic residue G176 to leucine, Tse6 catalytic residue D396 to alanine, Tse7 catalytic residues H229 and H230 to alanine, Tse8 catalytic residues S186 to alanine, in-frame deletion of *tse5*, short for DUEC | This study |
| | Δ*retS* Δ*tssB1* Δ*tssB2* Δ*tssB3* | T6SS-null mutant, short for T6SS$^{-}$ | This study |
| *Vibrio cholerae* V52 | *rhh* | Strain used for competition assay and Cre delivery assay | [3] |
| | *rhh* Δ*tssM* | Strain used for competition assay and Cre delivery assay | [3] |
| *Escherichia coli* | | | |
| T-Fast | | Strain used for cloning and gene expression | TIANGEN |
| MG1655 | | Strain used for competition assay | Lab stock |
| SM10 λpir | | Strain used for competition assay and Cre delivery assay, with streptomycin resistance | This study |

| Primers | Sequences (5'-3') | Descriptions |
|---|---|---|
| pPSV37-RBS-hifi-R | gggatccaggaggaaacgatg | Reverse primer to amplify pPSV37 vector |
| pPSV37-Tat-hifi-R | actgcggcgcaagcggcg | Reverse primer to amplify pPSV37 vector with Tat signal peptide |
| pPSV37-FLAG-hifi-F | gattacaaggacgacgatgacaagaagctttagcataaccccttggg | Forward primer to amplify pPSV7 vector with FLAG tag |

| | | |
|---|---|---|
| pPSV37-F | tgtgtggaattgtgagcggata | Forward confirmation primer of pPSV37 vector |
| pPSV37-R | tttgatcgcccttcccaaca | Reverse confirmation primer of pPSV37 vector |
| Cre-hifi-F | tccaatttactgaccgtacaccaa | Forward primer to amplify pPSV37 vector with Cre |
| Cre-pPSV-F | ggggatccaggaggaaacgatgtccaatttactgaccgtacacca | Forward primer to amplify Cre |
| Cre-pPSV-R | cacttgtcatcgtcgtccttgtaatcatcgccatcttccagcagg | Reverse primer to amplify Cre |
| VgrG1a-pPSV-RBS-F | ggggatccaggaggaaacgatgcaactgacccgcctggtcca | Forward primer to amplify VgrG1a |
| VgrG1a-pPSV-R | cacttgtcatcgtcgtccttgtaatctcccttcgctggtggtggaaacatcgcc tgtacctgg | Reverse primer to amplify VgrG1a |
| VgrG1a-Cre-hifi-R | ttggtgtacggtcagtaaattggatcccttcgctggtggtggtggaaacatcg cctgtacctgg | Reverse primer to amplify VgrG1a |
| VgrG1b-pPSV-RBS-F | ggggatccaggaggaaacgatggcacttgcgcaacag | Forward primer to amplify VgrG1b |
| VgrG1b-pPSV-R | cacttgtcatcgtcgtccttgtaatcgttctggaggatcttgcgtcc | Reverse primer to amplify VgrG1b |
| VgrG1b-Cre-hifi-R | ttggtgtacggtcagtaaattggagttctggaggatcttgcgtcc | Reverse primer to amplify VgrG1b |
| VgrG1c-pPSV-RBS-F | ggggatccaggaggaaacgatggctattggccagcctttcg | Forward primer to amplify VgrG1c |
| VgrG1c-pPSV-R | cacttgtcatcgtcgtccttgtaatcacagttgatatcgacattgggc | Reverse primer to amplify VgrG1c |
| VgrG1c-Cre-hifi-R | ttggtgtacggtcagtaaattggaacagttgatatcgacattgggc | Reverse primer to amplify VgrG1c |
| Tse6-281-pPSV-RBS-F | gggatccaggaggaaacgatggatgcgcaagccgccgcccgact | Forward primer to amplify 1-281aa of Tse6 |
| Tse6-281-pPSV-R | cacttgtcatcgtcgtccttgtaatcgtcgaggttgccggaacg | Reverse primer to amplify 1-281aa of Tse6 |
| Tse6-281-Cre-hifi-R | ttggtgtacggtcagtaaattggagtcgaggttgccggaacg | Reverse primer to amplify 1-281aa of Tse6 |
| Tse6-179-223-R | ccgaggcggtcgccgaggtcggtgtgcagccattcctcca | Reverse primer to amplify 179-223aa of Tse6 |
| pPSV-Tse6-NTMD-hifi-F | gacctcggcgaccgcctcgg | Forward primer to amplify 224-281aa of Tse6 |
| Tse6-62-pPSV-F | gggatccaggaggaaacgatgatgttccagatcgtcaagggc | Forward primer to amplify 62-223aa of Tse6 |
| Tse7-225-pPSV-RBS-F | ggggatccaggaggaaacgatggccaacgaggtgtacgc | Forward primer to amplify 1-225aa of Tse7 |
| Tse7-225-pPSV-R | cacttgtcatcgtcgtccttgtaatcctgcgggcagcaactgttc | Reverse primer to amplify 1-225aa of Tse7 |
| Tse7-225-Cre-hifi-R | ttggtgtacggtcagtaaattggactgcgggcagcaactgttc | Reverse primer to amplify 1-225aa of Tse7 |
| Tse5-1145-pPSV-RBS-F | ggggatccaggaggaaacgatgagcggcctaccggtttc | Forward primer to amplify 1-1145aa of Tse5 |
| Tse5-1145-pPSV-R | cacttgtcatcgtcgtccttgtaatccaggccgagcgggtcctgg | Reverse primer to amplify 1-1145aa of Tse5 |
| Tse5-1145-Cre-hifi-R | ttggtgtacggtcagtaaattggatcctcctcctgcggccgccaggccgag cgggtcctgg | Reverse primer to amplify 1-1145aa of Tse5 |

**References:**


1. Hersch, S. J., Lam, L. & Dong, T. G. Engineered type six secretion systems deliver active exogenous effectors and Cre recombinase. *mBio* **12**, e0111521 (2021).
2. Pei, T. T. *et al.* Intramolecular chaperone-mediated secretion of an Rhs effector toxin by a type VI secretion system. *Nat. Commun.* **11**, 1865 (2020).
3. Kamal, F. *et al.* Differential cellular response to translocated toxic effectors and physical penetration by the type VI secretion system. *Cell Rep.* **31**, 107766 (2020).
